# Influence of nanopillar arrays on fibroblast motility, adhesion and migration mechanisms

**DOI:** 10.1101/617001

**Authors:** Kai S. Beckwith, Sindre Ullmann, Jakob Vinje, Pawel Sikorski

## Abstract

Surfaces decorated with high aspect ratio nanostructures are a promising tool to study cellular processes and design novel devices to control cellular behaviour, perform intracellular sensing or deliver effector molecules to cells in culture. However, little is known about the dynamics of cellular phenomenon such as adhesion, spreading and migration on such surfaces. In particular, how these are influenced by the surface properties. In this work, we investigate fibroblast behaviour on regular arrays of 1 micrometer high, polymer nanopillars with varying pillar to pillar distance (array pitch). NIH-3T3 fibroblasts spread on all arrays, and on contact with the substrate engulf nanopillars independently of the array pitch. As the cells start to spread, different behaviour is observed. On dense arrays which have the pitch equal or below 1 micrometer, cells are suspended on top of the nanopillars, making only sporadic contact with the glass support. Cells stay attached to the glass support and fully engulf nanopillars during spreading and migration on the sparse arrays which are characterized by a pitch of 2 micrometers and above. These alternate states have a profound effect on cell migration rates, which are strongly reduced on nanopillar sparse arrays. Dynamic actin puncta colocalize with nanopillars during cell spreading and migration. Strong membrane association with engulfed nanopillars might explain the reduced migration rates on sparse arrays. This work reveals several interesting phenomenon of dynamical cell behaviour on nanopillar arrays, and provides important perspectives on design and applications of high aspect ratio nanostructures.

## 1 Introduction

Cells on vertically aligned high aspect ratio nanostructures (VA-HARNs) has been an area of great research interest in recent years.^1^ Examples include surface-based delivery of molecules to cells aided by the nanostructures,^2–9^ intracellular electrical measurements,^10–12^ capture of circulating tumour cells,^13–15^ induction of stem cell phenotypes,^16,17^ intracellular sensing,^18–20^ site-specific cellular imaging^11,21,22^ or controlling the geometry of in vitro neuronal networks.^23^ Despite the range of applications, there is still much unknown about how cells are influenced by HARNs. High viability and reduced spreading area of adherent cells is often reported.^24^ Arrays of nanowires have been shown to inhibit fibroblast migration with longer nanowires having a stronger effect,^25^ and small patches of nanopillars inhibited neuronal cell migration.^26^ Reduced spreading of cells on high aspect ratio nanostructures arrays has been reported,^16,27^ as well as altered expression of genes related to the cytoskeleton and cell adhesion.^28,29^ It has also been demonstrated that the cell membrane is able to wrap tightly over high aspect ratio nanostructures while maintaining membrane integrity.^30–32^ Careful investigations of the membrane integrity in cardiomyocyte-like HL-1 cells and Human embryonic kidney cells (HEK 293) on a variety of nanostructures done by Dipalo *et al* showed that vertical nanostructures can spontaneously penetrate the cellular membrane only in rare cases and under specific conditions.^30^

Surface bound nanostructures including nanopillars were used to show that local membrane curvature can directly act as source of a biochemical signal for endocytic proteins.^33,34^ Positively curved membranes were shown to be clathrin-mediated endocytosis (CME) hot spots, as determined by preferential accumulation of CME-related proteins at these sites. Nanopillars were also applied for controlled probing of nuclear mechanical properties. ^35^ In that report, the mechanism of nuclear deformation was linked to adhesive actin patches associating with the nanopillars, pulling the nucleus down, as well as a stress-fiber linked actin “tent” at the apical membrane exerting a downwards force.

Depending on the HARN array pitch, two distinct cell adhesion states have been reported: cells can be suspended on top of the nanostructures in a “bed of nails” effect, or adhere to the substrate with the HARNs protruding into, and even through the cell body with the cell plasma membrane wrapped around nanostructures.^16,24,27,36^ The critical array pitch for each state has been theoretically indicated to depend on nanostructure geometry, and has been experimentally shown to depend on surface chemistry, which influences cell-substrate adhesion.^36^ We have previously shown that on 1µm high nanopillar arrays, interim and dynamic states are also possible, where only smaller regions of the cell maintains substrate contact at a given time.^27^ Focal adhesions on nanowire arrays have been shown to be somewhat upregulated in one report,^37^ however they localized mainly on the substrate between nanowires. Carefully selected nanostructures have the potential to facilitate targeting and high throughput studies of certain molecular events in living cells. This can for example include endocytosis, formation of adhesion points, receptor clustering, dynamic of the cytoskeleton, cell membrane mechanics, etc. Much is still unknown about what type of structures are needed to activate certain biochemical signals and how these could be used in both screening and fundamental cell biology research. This is because most reported studies are performed on static, fixed cells. Substrates used for HARN are typically opaque, limiting possibilities of detailed dynamic studies of cell-nanostructure interactions with optical microscopy. Throughput of employed nanofabrication processes is often a significant bottleneck as well, as cell studies require many samples with relatively large area (~cm^2^). To better understand the cell responses to VA-HARNs and how they might be manipulated, we investigate the dynamics of NIH-3T3 fibroblasts adhesion, spreading and migration on arrays of high aspect ratio polymer nanopillars as a function of array pitch. Used cells are NIH-3T3 fibroblast stably transfected with LifeAct-mNeonGreen, ^38^ and PH-PLCd1-mScarlet^39,40^ which allow for visualization of F-actin and membrane respectively. LifeAct is a 17-amino-acid long peptide, that stains filamentous actin structures in eukaryotic cells and tissues.^38^ We employ nanopillar arrays fabricated with electron beam lithography (EBL) directly on microscopy-grade glass. The nanopillars are made from the stable, stiff and cell compatible polymer SU-8.^27^ We show that lamellipodium-induced membrane and cytoskeleton configuration around nanopillars strongly depends on pillar spacing on the substrate (array pitch). Lamellipodium-induced cell adhesion occurs immediately once the cells encounter the nanopillars during settling. Subsequent spreading and resulting cell adherence states are strongly influenced by the nanopillar pitch. During cell migration, cells form highly dynamic F-actin bundles at nanopillar sites, while focal adhesions form on the substrate between the nanopillars. Cell migration is reduced on sparse nanopillar arrays, but unchanged on denser arrays displaying the “bed of nails” effect. Our detailed dynamic studies shed new light on the range of mechanobiological interactions that may occur between cells and nanostructures.

## 2 Results

### 2.1 Nanopillar arrays

Nanopillar arrays were fabricated similar to our previously described approach,^27^ but with an 100keV Elionix GLS-100 EBL system, which allows to produce pillars with a more uniform tip/base diameter and with a much higher fabrication throughput (pattern writing speed >1 mm^2^/min). Figure 1 shows examples of fabricated nanopillar arrays illustrating uniform and reproducible nanopillar geometry. Nanopillar tip diameter is in the range of 90nm, and the diameter at the base is 130 nm for pillars 1µm in length used in this study. For fluorescence imaging, the nanopillars could be optionally doped with a fluorescent dye, for example oxazine-170 (far-red excitation and emission), as shown in Figure 1C. Hexagonal arrays of nanopillars with pitches of 0.75 - 10µm were made in mm^2^ areas on glass coverslips mounted into polystyrene cell culture dishes (35 mm dishes or 96-well plates). Nanopillar array pitches ≤ 1µm are considered dense, while pitches of ≥ 2µm are considered sparse. This setup for nanopillar arrays is ideal for high resolution microscopy of the interface between cells and nanostructures due to the possibility of imaging live cells in inverted optical microscope and with the use of low working distance, high numerical aperture objectives. Prior to investigating cell migration and attachment on the nanopillar arrays, we verify that cell membrane and actin filaments can be visualized using employed labelling strategy. Figure 2A and Figure 2B show micrographs of NIH-3T3 fibroblasts cultured on a glass substrate for 24h. PH-PLCd1-mScarlet label (orange) is concentrated on the plasma membrane, as well as on few intracellular vesicles (Figure 2A). LifeAct-mNeonGreen signal (green) clearly visualizes cytoskeletal F-actin architecture (Figure 2B).

**Figure 1:**
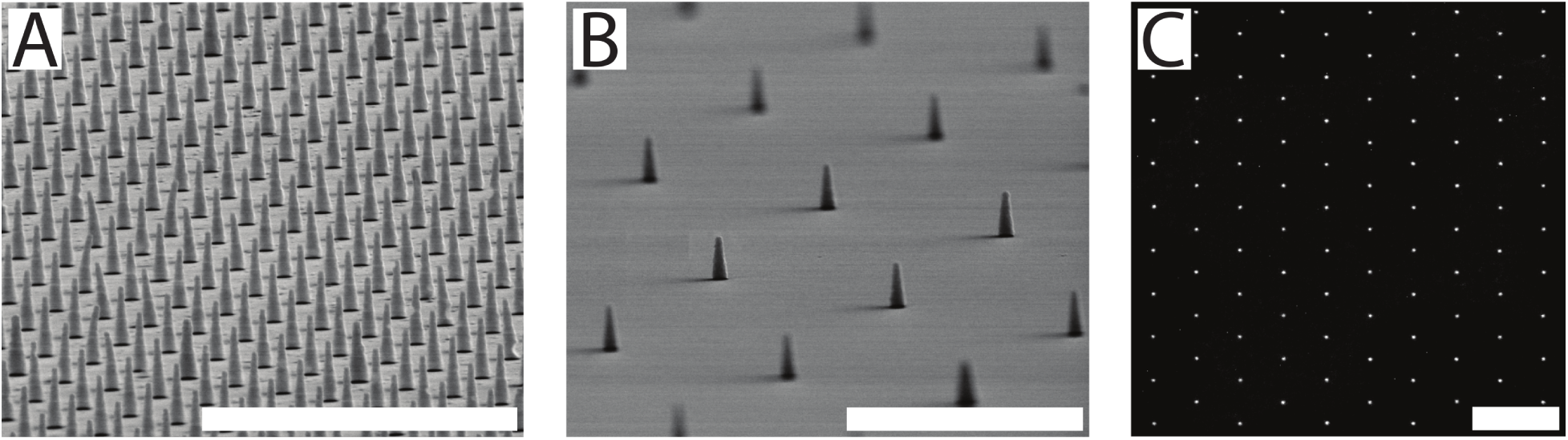
Overview of the nanopillar arrays used in this work. Tilted scanning electron micrographs of hexagonal SU-8 nanopillar arrays on glass cover slips with (A) 0.75µm and (B) 2µm pitch. (C) Confocal laser scanning microscopy image of 5µm pitch nanopillars array made from SU-8 doped with 100 µg/mL oxazine-170 dye. Scale bars 5µm in (A) and (B) and 10µm in (C).

**Figure 2:**
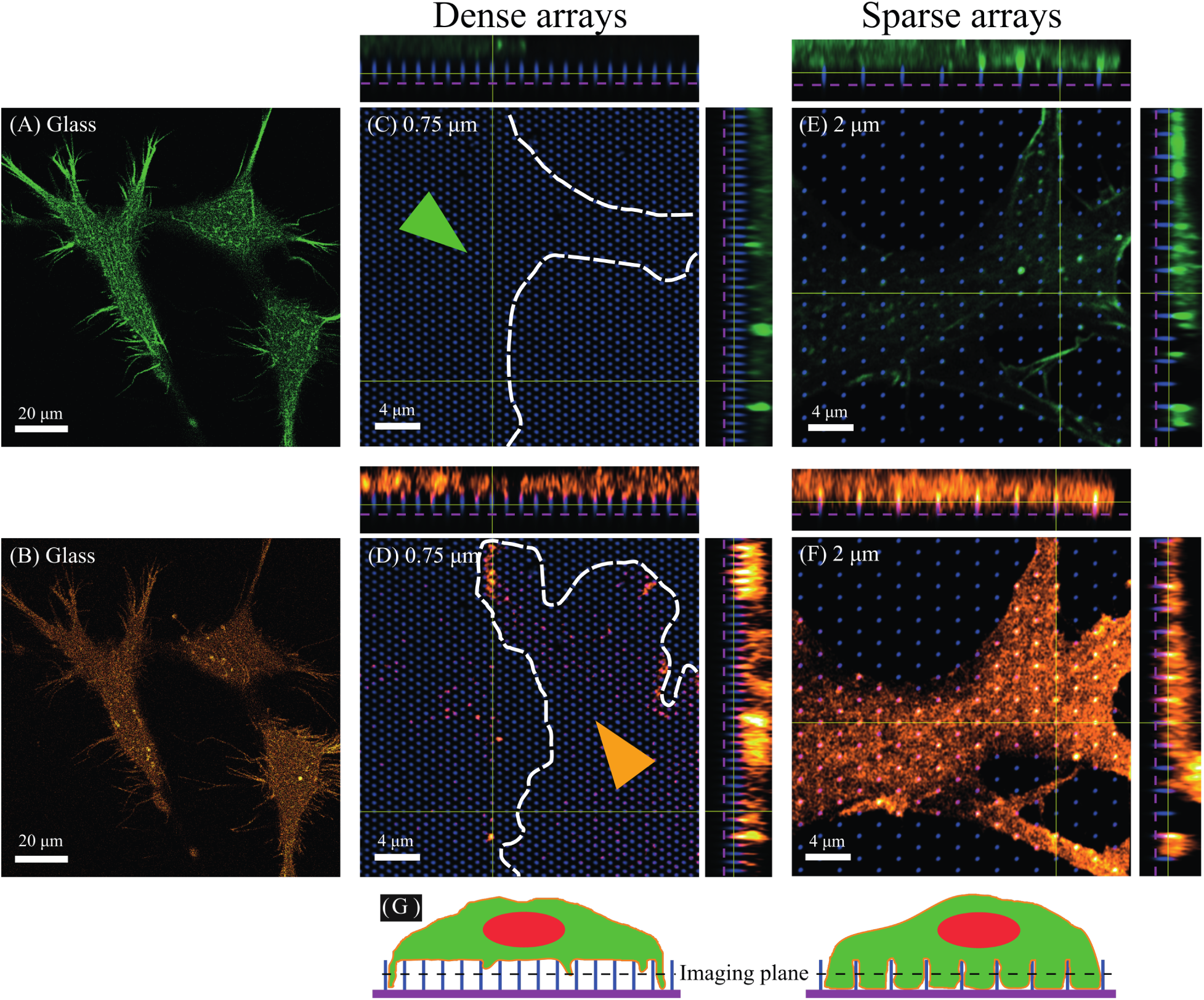
Initial characterization of NIH-3T3 fibroblasts expressing LifeAct-mNeonGreen (green) and PH-PLCd1-mScarlet (orange) localizing to actin and plasma membrane respectively. Single plane images acquired right above the glass substrate of the F-actin signal (A) and membrane signal (B) recorded for cells cultured on a glass substrate for 24h. High resolution Airyscan z-stacks with orthogonal projections of cells on arrays with a pitch of 0.75µm (C, D) and 2µm (E, F). Central, single plane images are taken at 50% of the pillar height. Dashed magenta lines in the side panels indicate the approximate location of the coverslip. Green arrow (C) and orange arrow (D) indicate position of the two cell bodies enclosed by white dashed lines. For the dense array no F-actin (C) and only fragmented membrane (D) signal is observed at the 50% height (500nm), indicating that the cell body is suspended on the top of the array. For the sparse arrays actin (E) and membrane (F) signals are observed for the whole cell, indicating that cells are in contact with the glass substrate and wrap around nanopillars. For the sparse arrays, membrane signal co-localizes with the signal from the pillars (see as pink colour). (G) Schematic of the two states observed on dense and sparse arrays.

Membrane and actin configuration for NIH-3T3 fibroblasts expressing LifeAct-mNeonGreen (green) and PH-PLCd1-mScarlet (orange) on dense and sparse naopillar arrays are presented in Figure 2C-F. The figure shows single confocal plane images taken at around 50% pillar height from the glass support (500nm), as well as *xz* and *yz* projections from corresponding *z*-stacks (side panels). Fluorescent signal from the Oxazine-170 dye doped SU-8 is shown in blue. Membrane signal co-localizes with the pillar signal along the pillar length for sparse arrays, seen as a pink color in both *xy*, *xz* and *yz* images (Figure 2F). As previously described, the signal enhancement in the membrane channel is due to the membrane wrapping around the nanopillars. ^27^ At some pillar locations F-actin signal is also clearly enhanced (F-actin puncta visible as bright green colour in the side panels, Figure 2E). For dense arrays, the plasma membrane is associated with pillars at the top of the pillar surface (Figure 2D, side panels). Some membrane wrapped around pillars at cell periphery, where cells make contact with the glass support, is also observed (Figure 2D, side panels). For these arrays, F-actin signal is only observed above the top surface of the pillar array, again indicating that most of the cell body is suspended on the top of nanopillar array (Figure 2C, side panels).

These results are corroborated by live-cell TIRF microscopy using the fluorescence signal from the cell membrane of NIH-3T3 fibroblasts expressing PH-PLCd1-mScarlet (Figure 3). For the glass control (Figure 3A), strong fluorescent signal is observed both in widefield fluorescence mode and in TIRF mode, indicating that the cell membrane is in immediate proximity to the glass substrate. The situation is different for cells on 0.75 and 1µm pitch nanopillar arrays (Figure 3B and Figure 3C), where strong signal is observed in the EPI mode, but only small patches of cell membrane are visible in the TIRF mode. These patches located close to the cell periphery and indicate that the cell membrane is in contact with the substrate only in these areas. The rest of the cell body is suspended on top of the nanopillar array. For sparse arrays, illustrated here by TIRF data for nanopillar spacing of 2µm, a larger area of the cell membrane is in contact with the glass substrate, indicating that the cell is wrapping around the pillars to a larger extent (Figure 3D).

**Figure 3:**
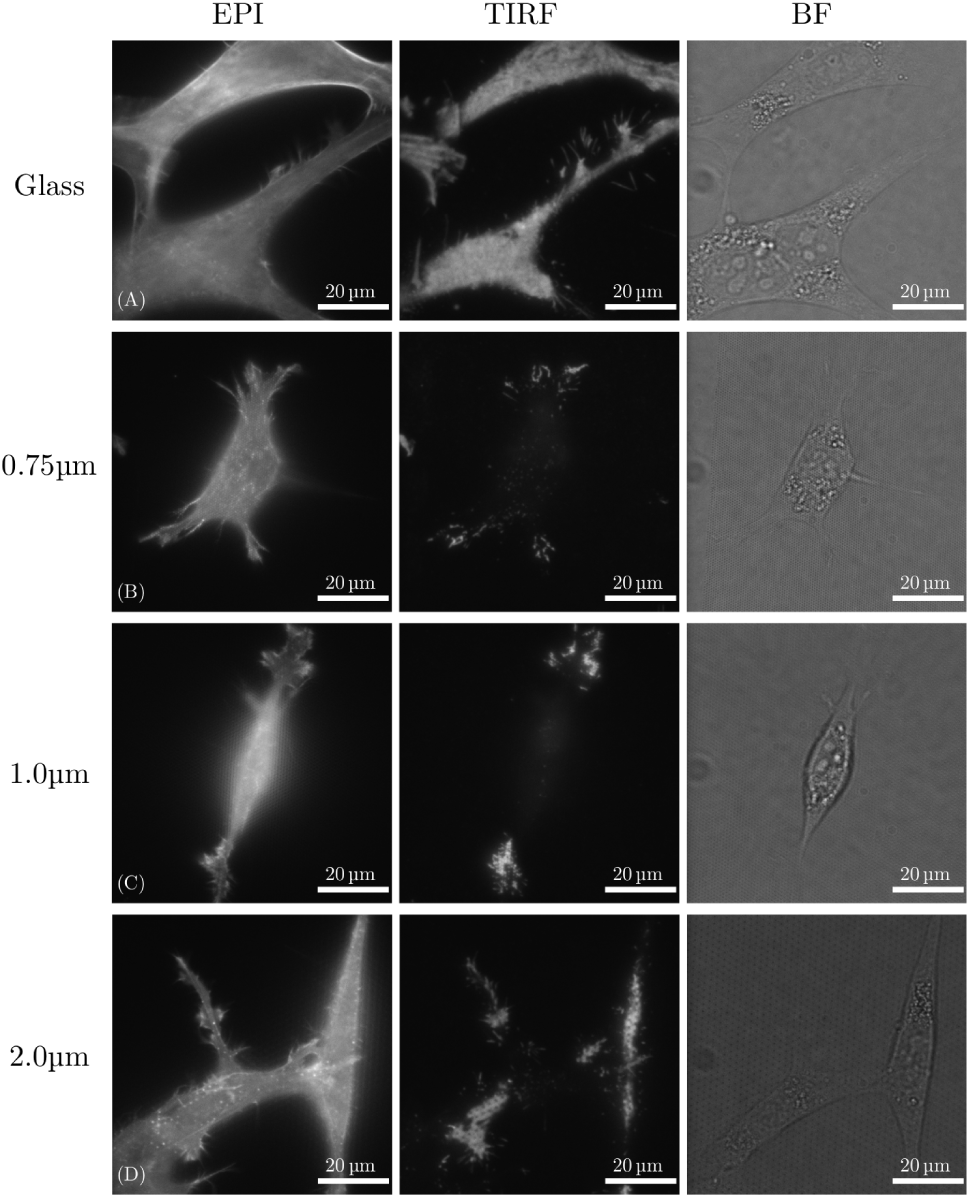
TIRF Data for NIH-3T3 fibroblasts 24h after seeding expressing PH-PLCd1-mScarlet localizing to the plasma membrane. Cells seeded on glass (A) give high signal from the membrane both in EPI fluorescence and in TIRF modes, indicating that the membrane is in contact with the glass surface. No TIRF signal from the cell bodies for cells on pillars with 0.75 and 1µm spacing (B and C) indicate that the membrane is not in contact with the glass except at the periphery of the cells. This contact is partially re-established on surfaces with larger pillar spacing, e.g. 2µm pitch arrays (D).

### 2.2 Fibroblast spreading on nanopillar arrays

In addition to describing cell conformation on various nanopillar arrays, we have used live cell microscopy to study the dynamics of initial cell attachment and settling of NIH-3T3 cells (Figure 4). Membrane and F-actin dynamics are further presented in Supplementary Movies 1 to 3. Flat, glass areas at the edge of the nanopillar arrays are shown to allow direct comparison of cell behaviour on these two surfaces. Cells were seeded onto nanopillar arrays after trypsination and imaged for *t* > 3h. On dense arrays, once making the initial contact with the substrate, the cells are able to engulf nanopillars and explore the glass surface (indicated cells in Figure 4A). However as soon as they start to spread, this contact is lost and the cells stay on the top of the arrays and only explore the glass surface at the cell periphery (Figure 4A and MovieS01). Moreover cells settling on the glass surface lose contact with the glass substrate when they migrate onto the dense pillar array, attaining a floating confirmation with lamellipodium at the leading edge reaching down to the glass surface (MovieS01). On sparse arrays nanopillars are also engulfed early in the settling process (Figure 4B and MovieS02 and MovieS03). However, in contrast to the dense arrays, this engulfment is stable during cell spreading and migration. F-actin and membrane signals are enhanced at the location of the pillars through the mechanism described above already after the initial contact with the surface. We observe a highly dynamic situation, where the F-actin puncta are forming and disappearing at various locations in the cell as a function of time (see below for more detailed description of this process).

**Figure 4:**
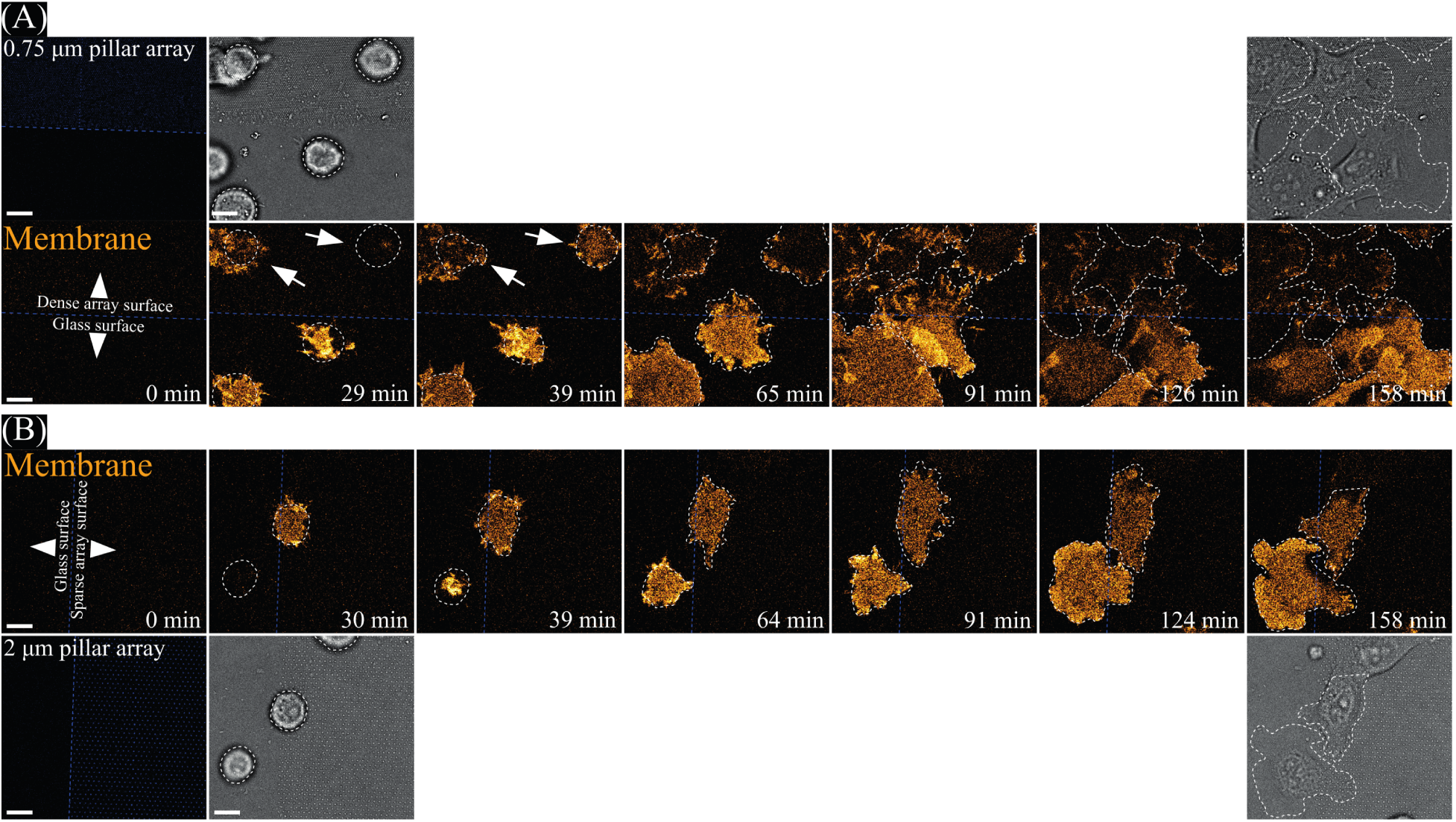
Confocal microscopy data recorded of NIH-3T3 fibroblasts expressing PH-PLCd1-mScarlet localizing to the plasma membrane settling onto dense (A, 0.75 µm spacing) and sparse (B, 2 µm spacing) pillar arrays. Images were acquired at approximately 50% pillar height. Blue dashed lines indicate borders between glass surface and pillar arrays, white dashed lines indicate approximate cell shapes based on bright field images. Cells were seeded with pillar samples positioned on the microscope such that imaging could commence immediately after seeding. For both dense and sparse arrays cells initially contact the glass coverslip both on pillar arrays and on glass (39min A and B). Note cells indicated with white arrows on the dense array after 29 and 39min also reach down to the glass coverslip. On dense arrays (A) the body of the cells subsequently pull up and membrane signal is mainly observed towards the edge of cells from 65min. On sparse arrays (B) the membrane signal underneath the cell bodies stay constant from the point of initial contact, indicating that cells stay in contact with the glass coverslip after initial contact both on and off pillar arrays (e.g. 64min). Scale bars 10 µm.

### 2.3 Fibroblast migration

Once the initial adhesion and spreading has occurred, fibroblasts start migrating. We performed large scale migration experiments by time-lapse imaging of NIH-3T3 cells on at least four 2.25mm^2^ areas of each nanopillar array for up to 14h. The fibroblasts were seeded at least 20h prior to imaging. Cell nuclei were labelled with a far-red fluorescent dye SiR-DNA and tracked (Figure 5A) using ImarisTrack (see Material and Methods). Flower plots of trajectories from cells migrating on glass and 2µm arrays are shown in Figure 5B-C. Due to unavoidable variations in cell seeding density, migration rates varied between independent migration experiments performed on the same substrates. Therefore, data from two large scale migration experiments were analyzed separately, pooling together data from individual wells recorded in parallel for each of the experiments (4-8 wells depending on the type of the array). Figure 5D presents a brief summary of the cell migration data. For arrays with 2µm pitch, migration data was recorded from a total area of 18mm^2^, a vary large area considering high resolution patterning. This area corresponds to 4.5×10^6^ pillars for 2µm arrays and to 32×10^6^ nanopillars for 0.75µm arrays.

**Figure 5:**
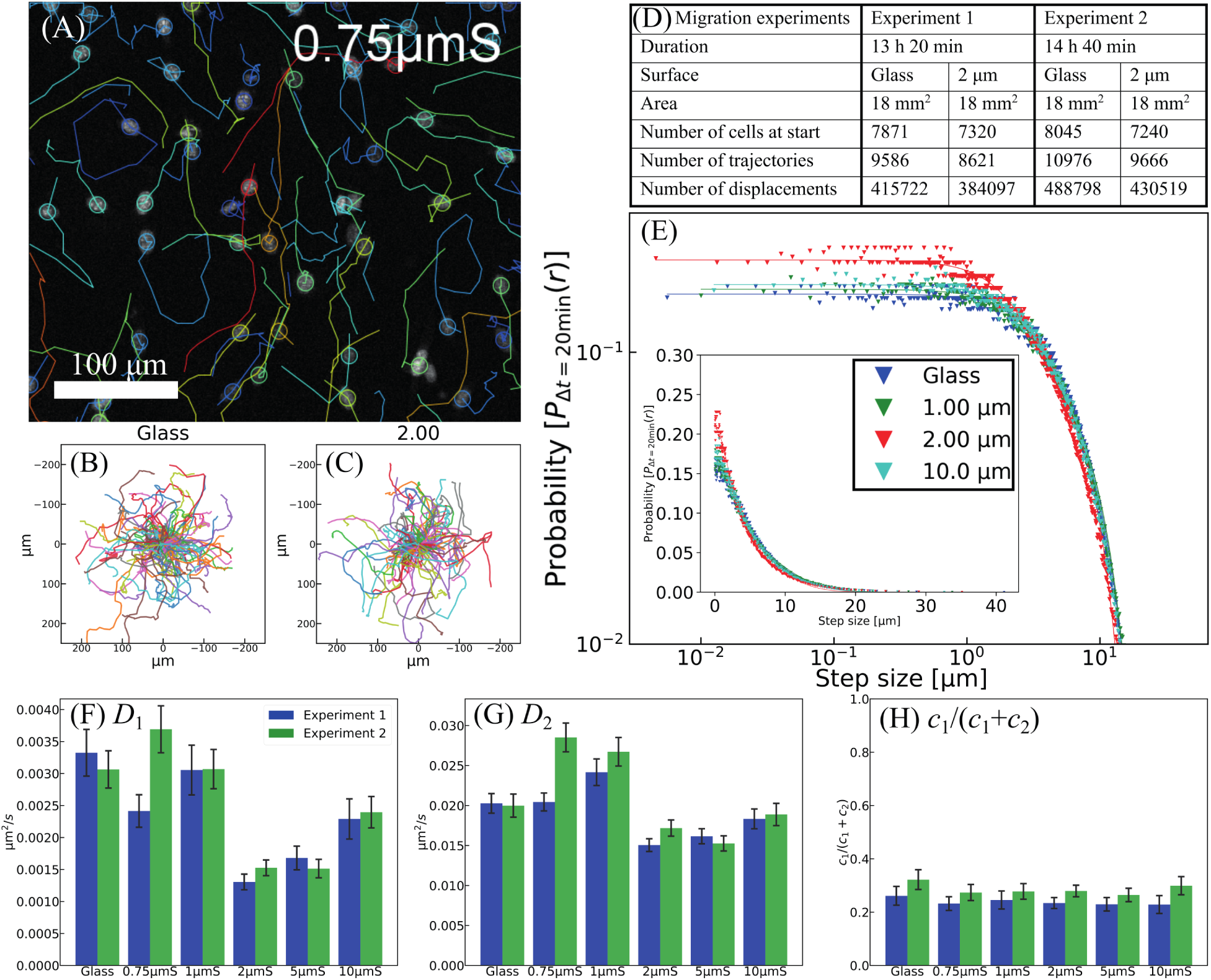
SiR-DNA-labelled nuclei of fibroblasts on nanopillar arrays and glass controls were imaged every 20min and tracked for up to 14h 40min. (A) Typical trajectories for tracked SiR-DNA stained nuclei in cells migrating on 0.75µm pitch arrays. In (B) and (C) representative sets of tracks are presented with a common origin. In (D) the surface area, number of cells, number of tracks and displacements analyzed for cells migrating on glass and 2µm pitch arrays are reported for the two presented experiments. For the four other pitch arrays (0.75, 1, 5 and 10µm) these numbers were halved due to the design of the pillar samples, glass and 2µm pitch arrays were present on all samples, while samples either had 0.75µm and 1µm, or 5µm and 10µm pitch arrays in addition. The cell displacements were analyzed by constructing a probability distribution of step sizes (filled triangles, on a log-log and inset linear plot) and fitting to a double Gaussian model (solid lines) presented in (D). Fitted parameters *D*_1_, *D*_2_ and *c*_1_ (weight coefficients depicted as *c*_1_/(*c*_1_ + *c*_2_)) from the Gaussian model are presented in (F-H). Error bars are 95% confidence intervals for each parameter extracted from the least squares fitting of the double Gaussian model to the migration data.

Due to large variations in the migration rates for individual cells, cell trajectories were analyzed by a method proposed by Banigan et.al.^41^ and described in detail in Materials and Methods. Briefly, for each data set, a probability density function of step sizes was generated and fitted to a double gaussian model. In this model, the heterogeneously migrating cell population is modelled as the sum of two (slow and fast migrating) randomly migrating cell populations with characteristic diffusion coefficients *D*_1_ and *D*_2_ as well as a weighting factors *c*_1_ and *c*_2_ for each population.

The resulting plots (Figure 5E) for Experiment 1, and determined values for diffusion coefficients for both experiments (Figure 5F and Figure 5G) demonstrate that cell migration is indeed inhibited on sparse nanopillar arrays compared to glass. Interestingly, the dependency on the nanopillar density is not directly proportional to nanopillar array pitch. On dense arrays (0.75-1µm) cell migration is not significantly hampered, but on sparse arrays migration is strongly reduced, with migration recovering on 10µm pitch. Comparing with the glass control, migration on the 2µm pitch substrates is reduced by 50% and 25% for the slow and fast migrating cell populations respectively. Based on not overlapping confidence intervals for data on glass and on 2µm pitch arrays, we can conclude that this difference is statistically significant with p-value which is at least *p* = 0.05.^42,43^ The slow population has a diffusion coefficient that is 7-10 times lower than the coefficient for the fast cell population on all substrates. The weighting factor, plotted as *c*_1_/(*c*_1_ + *c*_2_) in Figure 5H, reveal that about 25% of the analyzed displacements are modelled in the less mobile *D*_1_ population.

### 2.4 High resolution investigation of fibroblast migration

To gain insight into the processes occurring during cell migration, especially the origin of observed migration rate differences, live-cell confocal imaging was performed on migrating fibroblasts expressing LifeAct-mNeonGreen and PH-PLCd1-mScarlet. Selected time points from migrating fibroblasts on nanopillar arrays are shown in Figure 6. On dense nanopillar arrays, fast dynamics of densely spaced F-actin puncta is observed (see inserts in Figure 6A and Supplementary MovieS04 - MovieS08). Both F-actin and membrane signal are more intense close to cell periphery, indicating that the cells are close to the glass substrate in these regions and wrap along the full length of the pillars. Only a small enhancement of the membrane signal at the locations of nanopillars is observed (Figure 6B). This indicates, that apart from the peripheral regions, only small and dynamic membrane indentations are formed for cells on dense arrays. On spares arrays, the membrane signal is highly enhanced and stable (Figure 6D), indicating that the cell membrane wraps along the full length of the nanopillars. The cell body maintain a stable contact with the glass support (see also Supplementary MovieS09 - MovieS12). On sparse arrays membrane at the trailing edge of the migrating cell appears to remain associated with the nanopillars, which could be one reason for reduction in cell migration on sparse arrays. In contrast to the membrane signal (Figure 6D) which is uniform and stable, F-actin puncta seen in Figure 6C are highly dynamic. This highlights transient recruitment and dis-assembly of F-actin at nanopillars locations.

**Figure 6:**
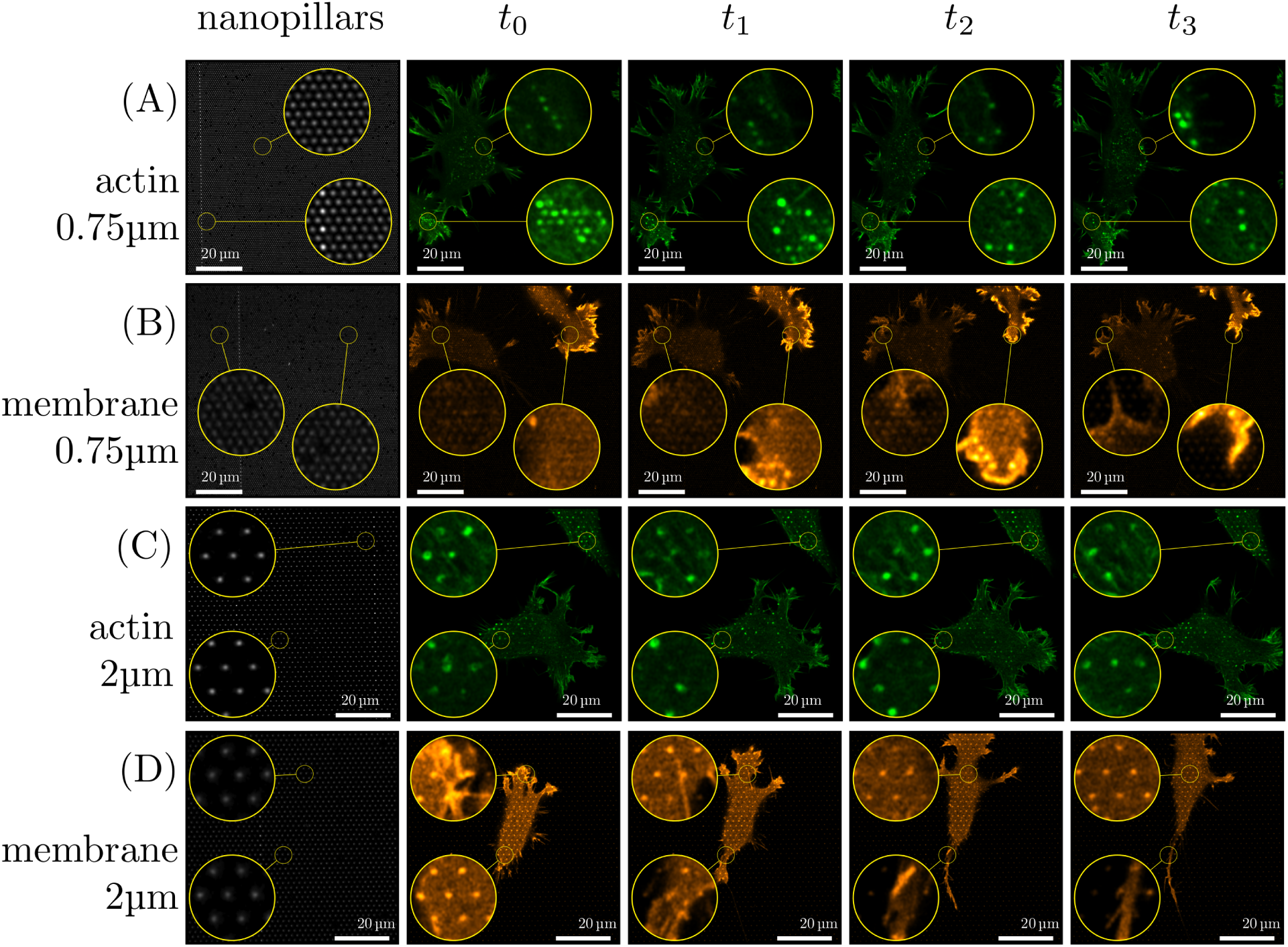
Migrating fibroblasts on nanopillar arrays. Sequence of images illustrating dynamical changes in F-actin and membrane configuration in cells migrating on dense and sparse nanopillar arrays. Circular inserts show a magnified view of the same sample region for the image sequence in each row. (A) highly dynamical F-actin puncta are observed on dense arrays, while the signal originating from slight membrane indentations on the nanopillars is more stable (B). For sparse arrays, F-actin puncta associate with the nanopillars and can sometimes be observed as rings and are also highly dynamic. Membrane signal on sparse arrays is constant and uniform for all nanopillar in contact with the cell (D). Time intervals (A) 2.5min (B) 5min (C) 5min (D) 10min.

To gain a better insight into F-actin assembly dis-assembly dynamics at nanopillars, time-lapses with increased frame rate were acquired by widefield microscopy. The F-actin dynamics is visualized by showing *xt* and *yt* projections at nanopillar positions indicated by white lines on the *xy* image (Figure 7 and Suplementary MovieS14 - MovieS16). Actin recruitment is transient on both dense and sparse arrays, but on the sparse arrays, F-actin is assembled for longer time periods and on a higher percentage of nanopillars. This indicates that F-actin association might be caused by the membrane curvature at the nanopillars, which is increased on the sparse arrays due to cell wrapping the full length of the pillars.

**Figure 7:**
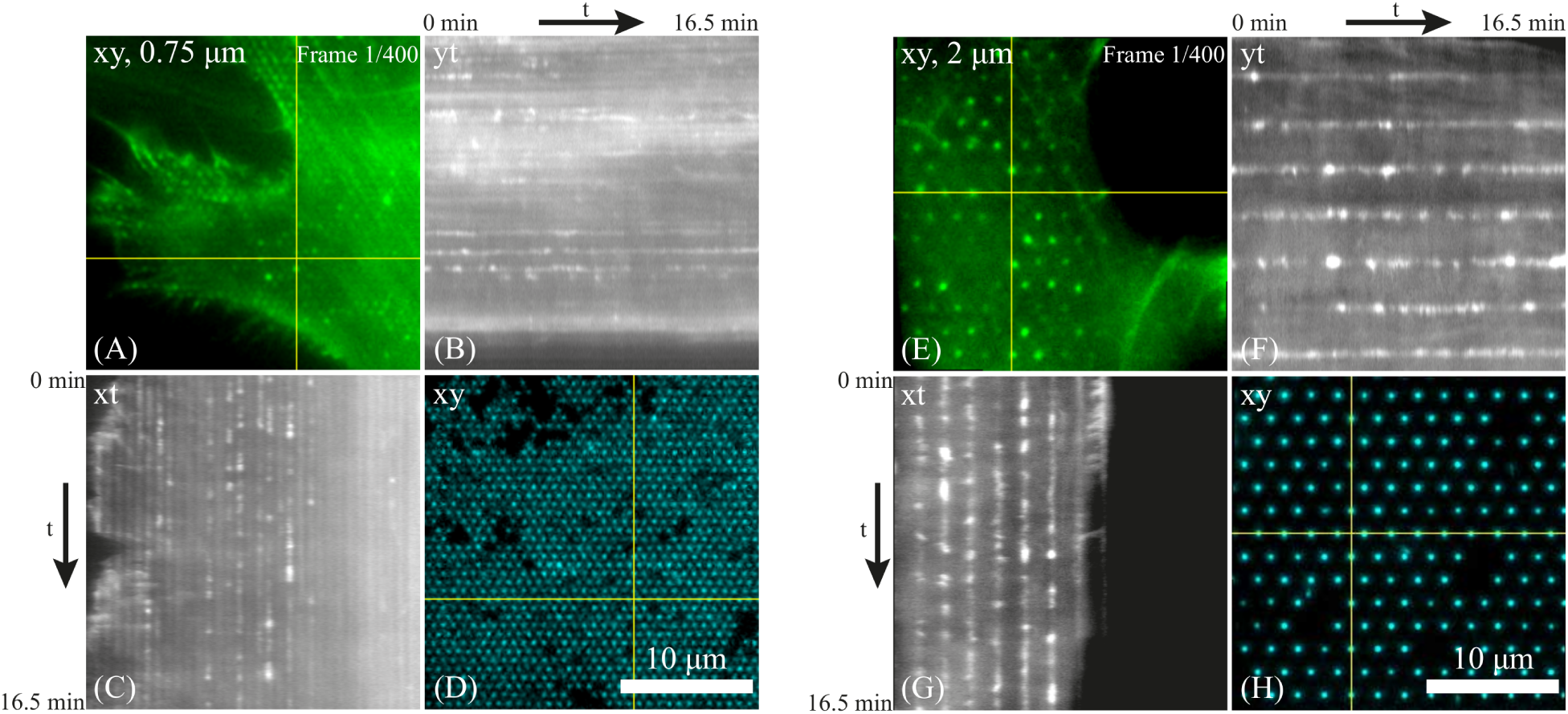
F-actin dynamics for NIH-3T3 fibroblasts expressing LifeAct-mNeonGreen recorded on (dense A-C) and sparse nanopillar arrays (E-G). Gray scale images show *xt* and *yt* projections along the indicated lines in the *xy* plane images shown for *t* = 0 (A,E) acquired at 2.5 s frame intervals. Corresponding nanopillar locations are shown in (D) and (H). Actin recruitment is transient on both dense and sparse arrays, but on the sparse arrays, actin is assembled for longer time periods and on a higher percentage of nanopillars. Also, intensity of the F-actin signal recorded for the sparse array is higher.

Finally, focal adhesion dynamics were investigated during cell spreading and migration (Figure 8). On both dense and sparse arrays actin puncta co-localizing with nanopillars was observed 30min after seeding, but focal adhesions had not yet formed. At 2h, actin puncta were still visible on both densities, and small, round focal adhesions formed between the nanopillars at the cell periphery, especially on the sparser arrays. At 4h, more elongated focal adhesions had formed on both sparse and dense arrays, and actin stress fibers were visible, especially on sparse arrays. Focal adhesions were formed mainly between nanopillars, but some signal overlapped with nanopillars as well. To investigate focal adhesion dynamics during cell migration, time-lapse microscopy was performed of fibroblast expressing talin-GFP. At the leading edge, new focal adhesions were observed to form mainly between the nanopillars, although some adhesions also appeared to form around or in very close vicinity to nanopillars. At the trailing edge focal adhesion dis-adhesion and retraction into the cell body was observed, despite the presence of nanopillars, although the focal adhesion appeared to circumvent the nanopillars during retraction. Thus, focal adhesion driven adhesion likely serves mainly to anchor the cells to the substrate, while actin and membrane interactions occur on the nanopillars.

**Figure 8:**
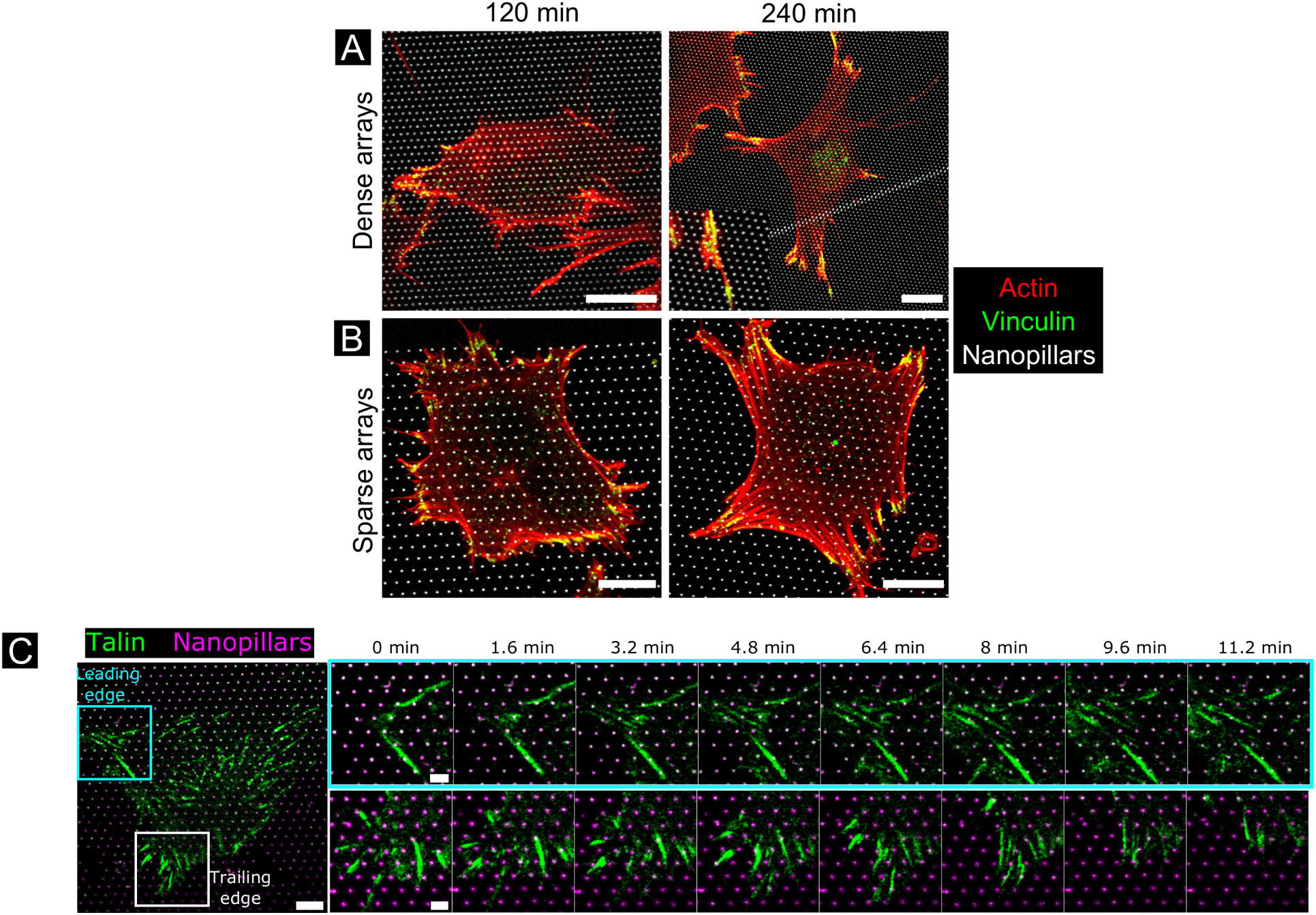
(A-B) Actin and vinculin organization in spreading fibroblasts. Fibroblasts were seeded for 2h and 4h before fixation and labelling Alexa488-phalloidin (red) and Alexa555-labelled vinculin (green) on dense arrays (1µm pitch) or sparse arrays (2µm pitch). Yellow color indicates actin-vinculin overlap. All scale bars are 10µm. (C) Excerpts from a time-lapse of a fibroblast transduced with talin-GFP (green) on a 2µm pitch nanopillar array (magenta), highlighting the formation of focal adhesions at the leading edge (cyan square) and trailing edge (white square) over a 11-min period. Scale bar overview image 10µm, excerpts 2µm.

## 3 Discussion

Cell adhesion, spreading and migration *in vitro* are highly studied processes, but have mainly been studied on flat substrates such as glass, or on gels or patterned surfaces. However, the geometry of the nanopillars introduce a new factor into the mechanobiology of cell adhesion and migration. To successfully manoeuvre the nanopillars, the main challenge for the cells appears to lie in shaping the membrane to conform to the nanopillars. Theoretical models of cell adhesion and membrane conformation on nanopillar arrays emphasis the balance between membrane bending and surface adhesion.^36,44,45^ Bending the cell membrane around each nanopillar requires energy to overcome the membrane bending stiffness, while adhesion to the surface of both the nanopillars and the substrate is energetically preferred for adherent cells used in this study. Thus, on dense nanopillar array pitches, the cells adopt a suspended state while on sparser arrays the cells adopt a conformal state, engulfing the nanopillars and adhering to the substrate. In this work, these two states were indeed observed in fully spread fibroblasts, with the transition occurring between 1µm and 2µm nanopillar array pitch. In addition, the detailed imaging performed here allowed several nuances to be observed.

During initial cell adhesion, nanopillar engulfment was observed regardless of nanopillar array pitch, which was not predicted in the theoretical models or observed experimentally before. The engulfment of the nanopillars likely has two interrelated contributing factors: membrane tension and actin polymerization forces. In general, depending on cell state and conformation, cells maintain significant membrane reserves stored as microscopic bends and buckles in the membrane.^46,47^ This could facilitate the initial engulfment of nanopillars regardless of array pitch. As the cell adhere and spread on the substrate, the membrane tension might increase, making the engulfment less preferred.

The time-lapse and selected time-point observations performed here additionally resolved intermediate states. Even in cells on dense arrays fully spread after 24h, small areas of cell membrane engulfing nanopillars were observed in cell peripheries. Likely, these protrusions are related to the second factor involved in membrane dynamics. Membrane movement at the leading edge (especially in lamellipodia) is driven by actin polymerization, resulting in a substantial force on the membrane, and is combined with enhanced adhesive complex formation.^48^ These interrelated phenomena of actin polymerization and focal adhesion formation were observed in this work at the leading edge of migrating cells and around the cell periphery of spread and spreading cells on all nanopillar arrays. It appears that in our system, on array pitches down to 0.75µm, actin polymerization and surface adhesions were sufficient to cause nanopillar engulfment at these locations, both initially, during cell adhesion and spreading, and later during cell migration. Once lamella replace lamellipodia, both actin polymerization and surface adhesion are diminished, resulting in suspension of the main cell body during spreading and migration on dense arrays. On sparse arrays the adhesive forces appear to remain strong enough to counter the increasing membrane tension, resulting in an adhered, nanopillar-engulfing state both during initial spreading and migration.

The distinction between these states has a strong influence on cell migration rates. Previous work with silicon nanowires and nanopillars has indeed shown strong reduction of migration in the sparse array regime, while less of a reduction was observed on dense arrays.^25,26,49,50^ The mechanisms for reduction in cell migration on sparse arrays are attributed to cytoskeletal and membrane entanglement by the nanostructures. Our results support these general descriptions, as in the sparse array regime both membrane engulfment and strong dynamic F-actin puncta association were observed. The requirement of continuous reorganization of the membrane and cytoskeleton in response to the moving cell likely causes a significant reduction of migration rates. A further effect of the nanopillars was observed where membrane residues were left behind on the nanopillars during retraction of the trailing edge. This indicates a strong adhesion between the membrane and nanopillars, sufficiently strong that the membrane does not release, but remains bound and forms membrane tethers. Strong membrane adhesion of cells to silicon nanowires has been reported,^28^ and extraction of membrane tethers is a well-known method of assessing membrane tension of cells.^51^ Such tethers are produced when an outward force is applied to a small area of the cell membrane and requires the loss of association between the cytoskeleton and cell membrane. Thus, the membrane binding and tether extraction at the trailing edge of migrating cells is a further hindrance for cell migration.

There are several opportunities for further investigations of cell migration on this platform. The exact mechanism of recruitment and function of the F-actin puncta or bundles around the nanopillars are not known. Using superresolution imaging we have previously shown that in HeLa cells these puncta are in fact rings or cylinders formed around the nanopillars. ^27^ In a recent report by Hanson *et. al.*, similar puncta and rings were observed and attributed to cell adhesion to nanopillars.^35^ As F-actin puncta were observed to be highly dynamic in this work, disappearing and appearing over a few min, it does not seem likely that these F-actin enrichments are solely involved in adhesion. Especially, as membrane engulfment of nanopillars was generally observed both with and without F-actin enrichment at the nanopillar site, this is not a continuous requirement for cell adhesion to nanopillars. We consider it likely that the dynamic F-actin enrichments are involved in the reorganization of the cell membrane as the cells migrate on nanopillar arrays, in addition to adherence processes. In a recent study, Mettlen et al showed that membrane curvature induced by using liposomes with various sizes, combined with PI(4,5)P_2_, and PI(3)P signalling are needed to trigger actin polarization.^52^ Some actin assemblies induced by membrane curvature on the sparse arrays (see Supplementary MovieS15) resembles actin comets induced in a cell-free assay for the role of phosphoinositide in actin polymerization.^52^ The role of nanopillar surface chemistry also remains to be investigated. Although geometry was attributed the leading role in reduced neuronal migration on nanopillars (Si, SiO_2_ and Pt were tested),^26^ it cannot be ruled out that the surface chemistry of SU-8 could play a role in the strong membrane interactions observed, and should be investigated further.

## 4 Conclusion

High resolution live cell imaging together with imaging of fixed cells and cell migration statistics reveal that nanopillars strongly influence both the quantitative and qualitative properties of fibroblast migration. From a fundamental perspective these results highlight the strong influence surface topography can have on cell function, inducing e.g. novel regimes of cell motility. From an applications perspective, the results demonstrate how important it is to investigate dynamic effects, as these reveal a nanostructure-cell interaction that changes both over time and even in different locations of the cell. The detailed interactions demonstrated here can have important implications for applications such as *in vivo* biointerfaces with nanowires or nanopillars, advanced *in vitro* applications such as neuronal guidance and network construction, stem cell differentiation and mechanobiology.

## 5 Experimental

### 5.1 Nanopillar fabrication

All chemicals and reagents were purchased from Sigma-Aldrich (Oslo, Norway) unless otherwise specified. SU-8 nanopillar arrays on glass were prepared as previously described.^27^ Briefly, 1µm thick SU-8 films were spin coated on 0.17mm (# 1.5) glass cover slips (Menzel-Gläser borosilicate glass) and soft-baked for 1min. Fluorescent SU-8 was made by including 100 µg/ml oxazine-170 in the SU-8 solution. Nanopillars were defined using an Elionix GLS-100 EBL-system. Samples were post-exposure baked for 3min and developed for 40s using mr-Dev 600 (Microchem, USA). The resulting nanopillars were 1µm high and had tip diameters in the range of 90 nm. All samples were treated with oxygen plasma before use.

### 5.2 Cell migration

NIH-3T3 cells (a kind gift from T. Sandal, NTNU) were maintained in DMEM with 10% FBS, 1% pen/strep, 1% MEM non-essential amino acids, 0.5% L-glutamine at 37°C. Seeding density was typically 25-30 thousand cells/cm^2^. 20-24h after cell seeding and 30-60min before commencing imaging, the far-red fluorescent dye SiR-DNA (Spirochrome) was added to the cell medium at a concentration of 1 µM labelling the cell nuclei. The cells were kept in the medium together with SiR-DNA for the duration of the migration experiment. Cells were imaged at 37°C with 5% CO_2_ using a 10X objective in an EVOS FL Auto 2 microscope. Cells were cultured on 3×3 mm nanopillar arrays, containing four 1.5×1.5mm^2^ areas with either 0.75, 1, 2µm pitch and glass control, or 2, 5, 10µm pitch and glass control. Images were recorded with 20min intervals (Δ*t* = 1200s) for up to 14h 40min. In each experiment four wells containing the dense arrays and four wells containing sparse arrays were imaged simultaneously. Nuclei positions were identified and tracked using the Imaris application ImarisTrack (Bitplane). From each experiment approximately 5 thousand cell trajectories were analysed for 0.75, 1, 5 and 10µm pitch and approximately 9 - 11 thousand tracks for 2µm pitch, giving a total of about 200 and 450 thousand cell displacements respectively. Probability density functions of cell displacements were analysed and fit to a double Gaussian function as described by Banigan *et. al*.^41^ Probability density function of cell displacements was constructed by assigning a chosen number of displacements *m* to each bin (*m* = 700 was chosen by Banigan et.al^41^ and was used in this work as well). The position of each bin was the average step size of that bin, while each bin was assigned a weight 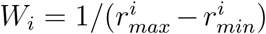 where 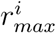 and 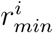 are the maximum and minimum displacements in that bin. This gives a normalized probability density function (which in our case summed to 2 as absolute values of all displacements were used). Note that we also pooled all *x* and *y* displacements. This probability density function was then fit to a double Gaussian of the form:

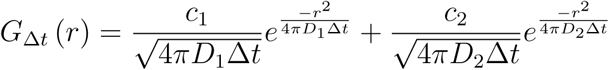

where *c*_1_ + *c*_2_ = 2 are the population weighting coefficients, *D*_1_ and *D*_2_ are the characteristic diffusion coefficient for each population, *r* is the cell displacement distance and Δt is the used frame rate. The double Guassian was fit to the probability density functions for each sample using least squares fitting, and 95% confidence intervals for each parameter were extracted as error estimates. Data was analysed using custom Python scripts and the `lmfit-py` package for fitting.^53^

### 5.3 High resolution cell imaging and immunofluorescence

NIH-3T3 cells were transduced with LifeAct-mNeonGreen targeting actin and the PH domain of PLC*δ*1 fused to mScarlet targeting the plasma membrane and stably selected on 10µg/mL blasticidine and 1µg/mL puromycin. The cells were imaged from 0-24h after seeding or during seeding after trypsination at 37°C and 5% CO_2_ with Zeiss LSM 800 Airyscan (Axiovert 200M inverted) CLS-microscope with a C-Apochromat 63x/1.15 W objective (Figure 6) or Zeiss LSM 880 Airyscan AxioObserver with a LD LCI Plan-Apochromat 40x/1.2W objective (Figure 2C-F) or Leica TCS SP8 HC PL APO CS2 63x/1.20W objective Figure 2A-C and Figure 4 or Zeiss Laser TIRF 3 with an alpha Plan-Apochromat 100x/1.46 Oil objective in both EPI fluorescence and TIRF mode (Figure 3 and Figure 7). NIH-3T3 cells were also transduced with BacMam 2.0 CellLights Talin-GFP (ThermoFisher) targeting focal adhesion and imaged at 37°C in Leibovitz’s L-15 Medium using a 63X 1.4 oil objective on a Leica SP8 microscope (Figure 8C). Cells were prepared for immunofluorescence using a protocol described by Whelan et al^54^ with some modifications. Briefly, cells were fixed in 4% PFA for 2.5min with no prior rinsing, washed in PBS, permeabilized in 0.1% triton x-100 in PBS for 5min, washed, blocked in 1% BSA in PBS for 15min, incubated with anti-vinculin (1:200) for 1h, washed, incubated with Alexa555 secondary antibody (1:500) for 1h, washed, and finally incubated with Alexa488-phalloidin for 30min. Fixed cells were imaged in PBS using a 63X 1.4 oil objective on a Leica SP8 microscope (Figure 8A-B) with system-optimized voxel sizes (typically 70 nm in *xy*).

## Acknowledgement

The authors would like to acknowledge Rasmus Schanke for initial experiments. The Research Council of Norway is acknowledged for the support to the Norwegian Micro- and Nano-Fabrication Facility (NorFab, grant number 245963/F50). NTNU is acknowledged for the financial support for JV through Nano@NTNU enabling technologies program and SU through NTNU Biotechnology enabling technologies program. This work was also partly supported by the Research Council of Norway through its Centres of Excellence funding scheme, project number 223255/F50.

## Supporting Information Available

### 5.4 Supplementary Data

**Table 1:** Values of fitted parameters *D*_1_, *D*_2_ and *c*_1_ weight coefficients from the double Gaussian model used to analyze the migration data (see Materials and Moethods). Weight coefficient *c*_2_ for the second population of cells can be calculated from: *c*_2_ = 2 − *c*_1_. Error bars (±) are 95% confidence intervals for each parameter extracted from the least squares fitting of the double Gaussian model to the migration data.

|  | Experiment 1 |  |  | Experiment 2 |  |  |
| --- | --- | --- | --- | --- | --- | --- |
| | $D_1$ [$\mu\text{m}^2 \text{s}^{-1}$]<br>$\times 10^{-3}$ | $D_2$ [$\mu\text{m}^2 \text{s}^{-1}$]<br>$\times 10^{-3}$ | $c_1$ | $D_1$ [$\mu\text{m}^2 \text{s}^{-1}$]<br>$\times 10^{-3}$ | $D_1$ [$\mu\text{m}^2 \text{s}^{-1}$]<br>$\times 10^{-3}$ | $c_1$ |
| Glass | $3.29 \pm 0.37$ | $20.0 \pm 1.2$ | $0.52 \pm 0.07$ | $3.07 \pm 0.29$ | $20.0 \pm 1.5$ | $0.64 \pm 0.07$ |
| 0.75 $\mu\text{m}$ | $2.42 \pm 0.26$ | $20.4 \pm 1.1$ | $0.46 \pm 0.05$ | $3.65 \pm 0.37$ | $28.0 \pm 1.7$ | $0.54 \pm 0.06$ |
| 1 $\mu\text{m}$ | $3.00 \pm 0.39$ | $23.6 \pm 1.6$ | $0.48 \pm 0.07$ | $3.07 \pm 0.31$ | $26.7 \pm 1.8$ | $0.56 \pm 0.06$ |
| 2 $\mu\text{m}$ | $1.31 \pm 0.13$ | $15.1 \pm 0.8$ | $0.47 \pm 0.04$ | $1.53 \pm 0.13$ | $17.2 \pm 1.0$ | $0.56 \pm 0.04$ |
| 5 $\mu\text{m}$ | $1.68 \pm 0.18$ | $16.2 \pm 0.9$ | $0.46 \pm 0.05$ | $1.52 \pm 0.14$ | $15.3 \pm 1.0$ | $0.53 \pm 0.05$ |
| 10 $\mu\text{m}$ | $2.29 \pm 0.32$ | $18.3 \pm 1.2$ | $0.46 \pm 0.07$ | $2.37 \pm 0.24$ | $18.5 \pm 1.3$ | $0.59 \pm 0.07$ |

### Supplementary Movies

- Examples of cells settling on dense and sparse nanopillar arrays. In all movies, NIH-3T3 fibroblasts expressing LifeAct-mNeonGreen (green) and PH-PLCd1-mScarlet (orange) were used and the cells were imaged for around 4h, capturing the initial cell attachment and spreading on various arrays. Nanopillar arrays are visible in the bright field images and in the oxazine-170 channel (blue). In some movies, pillars cover only part of the imaged sample area, so the cell behaviour on glass can be directly compared with the behaviour on the pillar arrays.

– MovieS01.avi: array pitch: 0.75µm. Total time: 220min
– MovieS02.avi: array pitch: 2.0µm. Total time: 217min
– MovieS03.avi: array pitch: 2.0µm. Total time: 217min
- Examples of cells migration on dense and sparse nanopillar arrays. In all movies, NIH-3T3 fibroblasts expressing LifeAct-mNeonGreen (green) and PH-PLCd1-mScarlet (orange) were used and the cells were imaged at 2.5 to 5min intervals, capturing cell migration dynamics at the time scale of <1h.

– MovieS04.avi. Array pitch: 0.75µm; frame interval 2.5min. F-actin signal.
– MovieS05.avi. Array pitch: 0.75µm; frame interval 2.5min. F-actin signal.
– MovieS06.avi. Array pitch: 0.75µm; frame interval 5min. Membrane signal.
– MovieS07.avi. Array pitch: 0.75µm; frame interval 5min. Membrane signal.
– MovieS08.avi. Array pitch: 0.75µm; frame interval 5min. Membrane signal.
– MovieS09.avi. Array pitch: 2.0µm; frame interval 2.5min. F-actin signal.
– MovieS10.avi. Array pitch: 2.0µm; frame interval 2.5min. F-actin signal.
– MovieS11.avi. Array pitch: 2.0µm; frame interval 5min. Membrane signal.
– MovieS12.avi. Array pitch: 2.0µm; frame interval 5min. Membrane signal.
- Example of a migration data set. NIH-3T3 fibroblasts expressing LifeAct-mNeonGreen (green) were imaged for 14h at 37°C, 5% CO_2_ using a 10X objective of the EVOS FL Auto 2 microscope (Thermo Fisher Scientific).

– MovieS13.avi
- Examples of actin and membrane dynamics taken at high frame rate using epi fluorescence microscope. NIH-3T3 fibroblasts expressing LifeAct-mNeonGreen (green) and PH-PLCd1-mScarlet (orange) were imaged using Zeiss Laser TIRF 3 with an alpha Plan-Apochromat 100x/1.46 Oil objective in EPI fluorescence mode. For dense arrays, only actin signal was recorded, due to very weak membrane signal.

– MovieS14.avi. Array pitch: 0.75µm; frame interval 2.2s. F-actin signal.
– MovieS15.avi. Array pitch: 2.0µm; frame interval 2.5s. F-actin and membrane signal.
– MovieS16.avi. Array pitch: 2.0µm; frame interval 1.8s. F-actin and membrane signal.

